# Serotonin, Etonogestrel and breathing activity in murine Congenital Central Hypoventilation Syndrome

**DOI:** 10.1101/2022.04.19.488470

**Authors:** Alexis Casciato, Lola Bianchi, Fanny Joubert, Roman Delucenay-Clarke, Sandrine Parrot, Nélina Ramanantsoa, Eléonore Sizun, Boris Matrot, Christian Straus, Thomas Similowski, Florence Cayetanot, Laurence Bodineau

## Abstract

Congenital Central Hypoventilation Syndrome, a rare disease caused by *PHOX2B* mutation, is associated with absent or blunted CO_2_/H^+^ chemosensitivity due to the dysfunction of PHOX2B neurons of the retrotrapezoid nucleus. No pharmacological treatment is available. Clinical observations have reported non-systematic CO_2_/H^+^ chemosensitivity recovery under desogestrel. Here, we used a preclinical model of Congenital Central Hypoventilation Syndrome, the retrotrapezoid nucleus conditional *Phox2b* mutant mouse, to investigate whether etonogestrel, the active metabolite of desogestrel, led to a restoration of chemosensitivity by acting on serotonin neurons known to be sensitive to etonogestrel, or retrotrapezoid nucleus PHOX2B residual cells that persist despite the mutation. The effect of etonogestrel, alone or combined with serotonin drugs, on the respiratory rhythm of medullary-spinal cord preparations from *Phox2b* mutants and wildtype mice was analyzed under metabolic acidosis. c-FOS, serotonin and PHOX2B were immunodetected. Serotonin metabolic pathways were characterized by ultra-high-performance liquid chromatography. We observed etonogestrel restored chemosensitivity in *Phox2b* mutants in a non-systematic way. Histological differences between *Phox2b* mutants with restored chemosensitivity and others indicated greater activation of serotonin neurons of the *raphe obscurus* nucleus but no effect on retrotrapezoid nucleus PHOX2B residual cells. Pharmacology of serotonin systems modulated the respiratory effect of etonogestrel differently according to serotonin metabolic pathways. Etonogestrel induced a restoration of chemosensitivity in *Phox2b* mutants by acting on serotonin neurons. Our work thus highlights that the state of serotonin systems was critically important for the occurrence of an etonogestrel-restoration, an element to consider in potential therapeutic intervention in Congenital Central Hypoventilation Syndrome patients.

## INTRODUCTION

Congenital central hypoventilation syndrome (CCHS; also named Ondine’s curse) is a rare life-threatening sleep-related central hypoventilation (incidence 1/148,000–1/200,000 live births) associated with an absent or blunted respiratory response to hypercapnia (Trang et al., 2020). CCHS is associated with an autosomal dominant mutation in the paired-like homeobox 2B gene (*PHOX2B*) which encodes a highly conserved homeodomain transcription factor (Amiel et al., 2003, Amiel et al., 2009). *PHOX2B* is considered as the disease-defining gene in CCHS (Weese-Mayer et al., 2017). The most common mutations are the addition of trinucleotides encoding alanine in exon 3, *i*.*e*. polyalanine mutations, and 7-alanine expansion is the most frequent *(PHOX2B*^*27Ala/+*^) (Amiel et al., 2003, Amiel et al., 2009). Other rare mutations, grouped as non-polyalanine, correspond to missense, frameshift, nonsense and stop codon mutations (Amiel et al., 2009). Specific loss of PHOX2B cells in the retrotrapezoid nucleus (RTN) is the primary neuroanatomical defect observed in *Phox2b*^*27Ala/+*^ mice and respiratory defects in CCHS are at least partly attributed to loss or dysfunction of CO_2_/H^+^ chemosensory PHOX2B neurons of the RTN (Dubreuil et al., 2008, Ramanantsoa et al., 2011, Weese-Mayer et al., 2017, Guyenet and Bayliss, 2015). In the absence of effective curative treatment, patients are placed on assisted ventilation for life, at least when sleeping (Weese-Mayer et al., 2010), and insufficient ventilatory support can expose patients to neural damage and impair their quality of life (Harper et al., 2014, Weese-Mayer et al., 2017).

We previously reported that two adult women with CCHS recovered CO_2_/H^+^ chemosensitivity concomitantly with the administration of desogestrel for contraception (Straus et al., 2010). This suggests that desogestrel or its biologically active metabolite, the 3-ketodesogestrel (etonogestrel), could activate or over-activate the residual CO_2_/H^+^ chemosensitivity present in certain CCHS patients (Carroll et al., 2014), an effect which was not provoked by the upsurge of progesterone during pregnancy (Sritippayawan et al., 2002). This observation undermined the prevailing view that respiratory symptoms in CCHS are irreversible, and paved the way for treatment research. Unfortunately, the recovery of chemosensitivity under desogestrel was not confirmed in a third patient (Li, 2013), suggesting that effect in CCHS patients was contingent on the status of yet unknown respiratory pathways. Such a target could be medullary serotonin (5-HT) neurons, which we have recently shown to express the activation marker *c-Fos* under etonogestrel in rodent models free from CCHS (Joubert et al., 2016b, Loiseau et al., 2019). Another possible explanation could be that etonogestrel acts on PHOX2B residual cells by decreasing the level of toxic cellular mutant PHOX2B protein, as recently reported in neuroblastoma cell lines (Di Lascio et al., 2020, Cardani et al., 2018).

Bearing in mind that desogestrel can induce recovery of CO_2_/H^+^ chemosensitivity in CCHS patients (Straus et al., 2010) and assuming that respiratory defects in CCHS patients are at least partly attributed to alteration in RTN PHOX2B cells (Dubreuil et al., 2008, Ramanantsoa et al., 2011), we postulated first that etonogestrel would induce a restoration of CO_2_/H^+^ chemosensitivity in RTN conditional *Phox2b* mutant mice, and second that it would act either on 5-HT medullary neurons, on residual RTN PHOX2B cells (Ramanantsoa et al., 2011), or both. To validate our hypotheses, we combined pharmacological applications, electrophysiological recordings on *ex vivo* preparations, functional and phenotypic histology, and ultra-high-performance liquid chromatography.

## RESULTS

### Recovery of the CO_2_/H^+^ chemosensitivity by etonogestrel in *Phox2b* mutants

Preparations from *Phox2b* mutants displayed a much slowed respiratory-like rhythm in normal-pH, significantly lower than that of wildtype littermates (0.8 ± 0.1 cycles.min^-1^, n=98 *vs* 13.9 ± 0.8 cycles.min^-1^, n=115; p<0.0001). Under metabolic acidosis, wildtype littermates showed increased respiratory frequency (fr, +29%, p<0.001; **Figure 1, Figure 2B, 1I**). For all mutant preparations collected, there was no response to metabolic acidosis as previously described (Ramanantsoa et al., 2011), but some increased their respiratory-like rhythm and others not. As in similar conditions (Wei and Ramirez, 2019), we designated two populations defined by the ability (“acidosis-responder”) or failure (“acidosis-non-responder”) to increase their respiratory-like rhythm by at least 10%: 4 of the 23 preparations (17%) were thus considered as acidosis-responders (**Figure 2J**).

**Figure 1.**
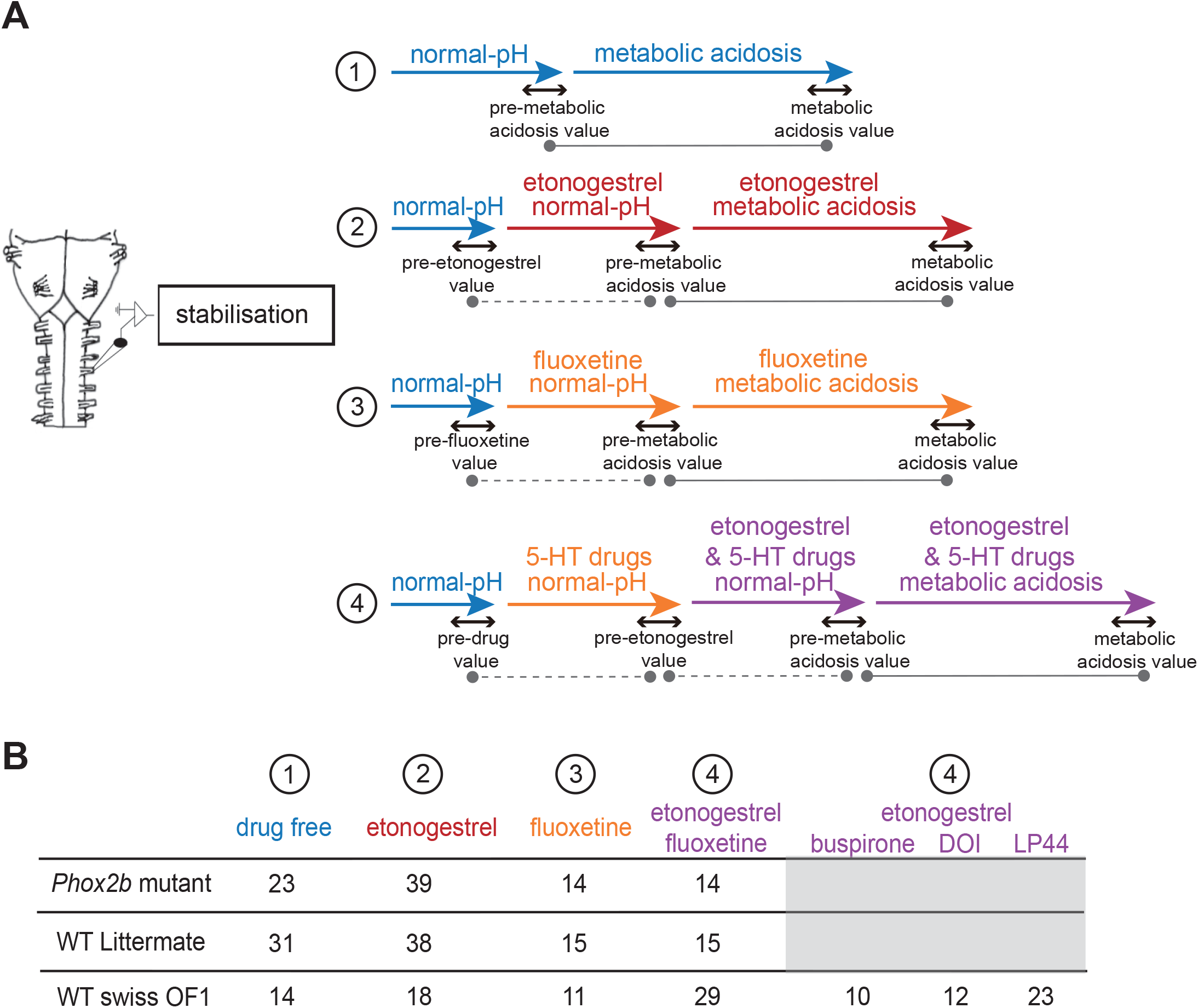
Schematic representation of the pharmacological protocols used. Flowchart of the protocols used in *ex vivo* medulla-spinal cord preparations from *Phox2b* mutants, wildtype littermates and wildtype Swiss OF1 mice (**A**). Briefly, after a period allowing fr to stabilize, preparations were subjected to one of four protocols: 1) preparations were exposed to drug-free normal-pH and metabolic acidosis conditions, 2) preparations were exposed to drug-free normal-pH followed by exposure to etonogestrel under normal-pH and then metabolic acidosis conditions, 3) preparations were exposed to drug-free normal-pH followed by exposure to fluoxetine under normal-pH and then metabolic acidosis conditions, 4) preparations were exposed to drug-free normal-pH followed by etonogestrel associated with fluoxetine under normal-pH and then metabolic acidosis conditions. Table indicating the number of *ex vivo* preparations used for each protocol in *Phox2b* mutants, wildtype littermates and wildtype Swiss OF1 mice (**B**). The dotted grey lines represent the comparison made to determine the influence of a drug in normal-pH. The grey lines represent the comparison made to determine the response to metabolic acidosis under drug-free or drug exposure conditions.

**Figure 2.**
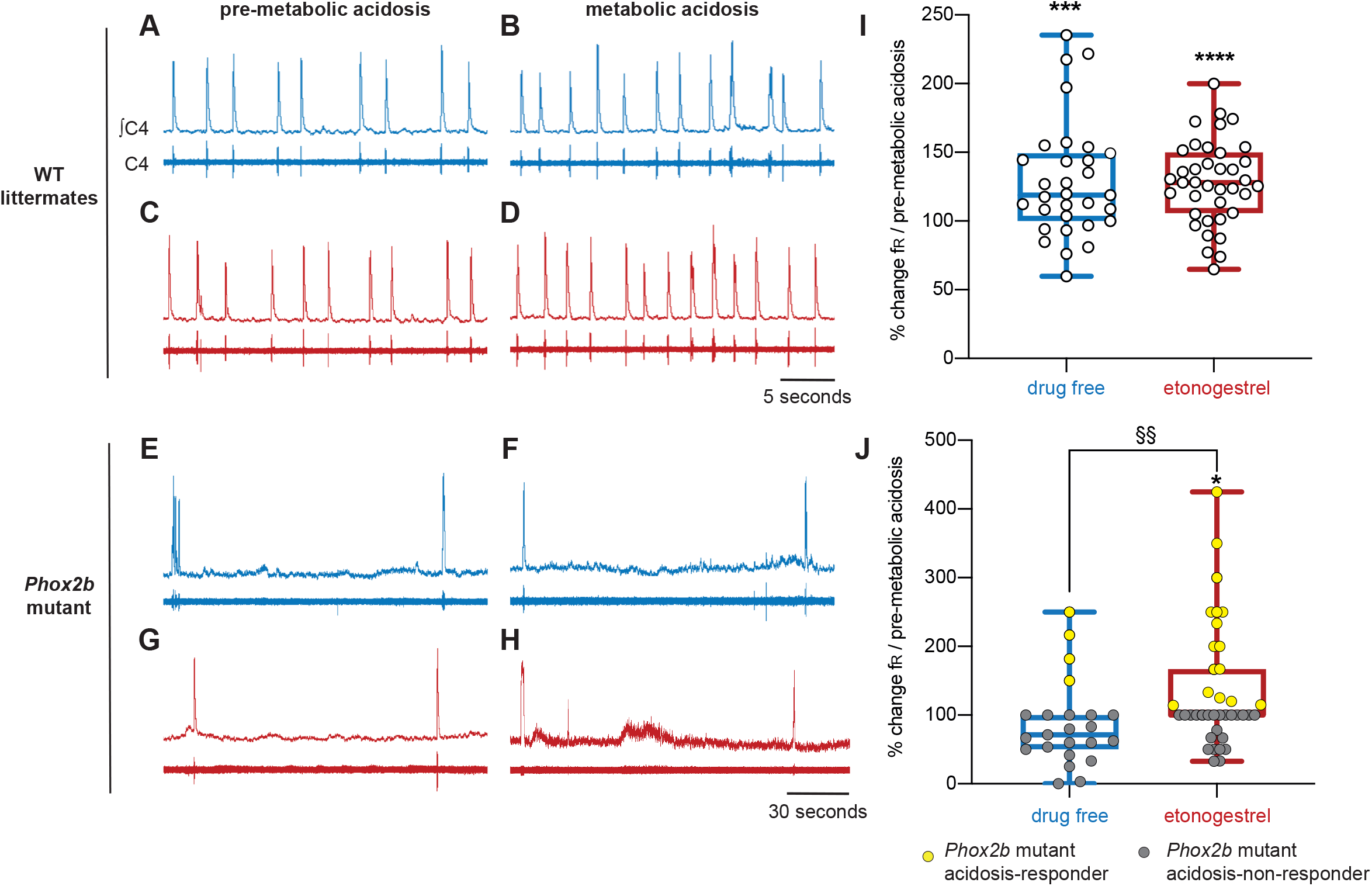
Effects of etonogestrel on central H^+^ chemosensitivity in *Phox2b* mutant mice. **A-H**, Individual phrenic activity traces (fourth cervical ventral nerve root, C4) and integrated C4 activity from medullary-spinal cord from wildtype littermates (**A**,**B**,**C**,**D**) and *Phox2b* mutant mice (**E**,**F**,**G**,**H**) preparations, under drug-free (**A**,**B**,**E**,**F**) or etonogestrel (**C**,**D**,**G**,**H;** 5.10^−2^ µM) exposure, in normal-pH (**A**,**C**,**E**,**G**; pre-metabolic acidosis) or metabolic acidosis conditions (**B**,**D**,**F**,**H**). **I-J**, Scatter plots with a superimposed box and whisker (median [Q1; Q3] and minimum and maximum values) showing the respiratory-like rhythm (respiratory frequency, fr) during the last five minutes of metabolic acidosis in percentage of pre-metabolic acidosis values under drug-free and etonogestrel exposure in wildtype littermates (**I**; n=31 and n=38 respectively) and *Phox2b* mutant mice (**J**; n=23 and n=39). Etonogestrel induced not only a restoration of the response to metabolic acidosis considering all preparations, but also an increase in the proportion of acidosis-responder (preparations displaying an increase in fr under metabolic acidosis compared to pre-metabolic acidosis values by at least 10%, p<0.05; Fisher’s exact test; **J**, yellow filled circles). * Indicates a significant difference in fr relative to pre-metabolic acidosis values (* p < 0.05, *** p < 0.001, **** p < 0.0001; paired t-test or Wilcoxon test). ^§^ Indicates a significant difference in *Phox2b* mutant mice between preparation exposed to metabolic acidosis with and without etonogestrel (^§§^ p < 0.01; Mann-Whitney test). ∫C4,integrated activity of C4 ventral nerve root; C4, electrical activity of C4 ventral nerve root, WT: wildtype.

Under etonogestrel, while the respiratory-like rhythm of *Phox2b* mutant preparations was not modified in normal-pH (p=0.33), it significantly increased (+35%; p<0.001) under metabolic acidosis conditions (**Figure 1, Figure 2H, 2J**). The proportion of acidosis-responder preparations was greater in presence of etonogestrel (16 of 39 preparations, 41% vs 17% without etonogestrel; p<0.05) than without the progestin (**Figure 2J**). In wildtype littermates preparations, etonogestrel led to a significant increase in fr in normal-pH compared to pre-etonogestrel values (16.4 ± 1.6 vs 13.0 ± 1.3 cycles.min^-1^, +29%, p=0.0003; **Figure 1, Figure 2C**), and did not modify the increase in respiratory-like rhythm induced by metabolic acidosis (+28%, 19.3 ± 1.4 cycles.min^-1^ vs 29% without etonogestrel; p<0.0001; **Figure 2D, 2I**).

### Difference in *c-Fos* expression under metabolic acidosis/etonogestrel between acidosis-responder and acidosis-non-responder *Phox2b* mutants

To identify the origin of the etonogestrel-associated recovery of H^+^ chemosensibility in *Phox2b* mutants, we compared the c-FOS number of cells in medullary respiratory areas between acidosis-responders (n=8) and acidosis-non-responders (n=10) (**Table 1, Figure 3**).

**Table 1.** c-FOS-positive number of cells in respiratory areas of the medulla oblongata in *Phox2b* mutant preparations under etonogestrel.

|  | acidosis-non-responder | acidosis-responder |
| --- | --- | --- |
| c/mNTS | 146.5 [121.1; 181.5] | 106.5 [83.0; 178.0] |
| cROb | 2.0 [1.0; 7.5] | 24.5 [9.5; 38] * |
| rROb | 5.5 [1.3; 9.0] | 10.5 [2.5; 50.8] |
| cRPa | 10.5 [7.0; 22.0] | 25.5 [10.0; 35.0] |
| rRPa | 74.5 [40.0; 102.0] | 57.0 [32.5; 152.0] |
| VLM | 263.5 [220.3; 318.5] | 449.0 [275.3; 497.3] |
| pf-RTN | 16.0 [12.8; 29.5] | 36.0 [29.8; 59.0] * |
| cRTN | 75.0 [62.9; 95.4] | 49.8 [34.0; 95.8] |
| pFRG | 19.0 [15.8; 35.8] | 29.0 [19.5; 45.5] |

Values as expressed as median number of total c-FOS-positive cells [Q1; Q3]
\* $P < 0.05$ , Responder *Phox2b* mutant values compared with Non-responder *Phox2b* mutant values
c/mNTS, commissural and median part of the solitary tract nucleus; cROb, caudal part of the nucleus of the raphe *obscurus* (from the pyramidal decussation to the rostral edge of the inferior olive); rROb; rostral part of the nucleus of the raphe *obscurus* (from the rostral edge of the inferior olives to the rostral edge of the facial nucleus); cRPa, caudal part of nucleus of the raphe *pallidus* (from the pyramidal decussation to the rostral edge of the inferior olive); rRPa, rostral part of nucleus of the raphe *pallidus* (from the rostral edge of the inferior olives to the rostral edge of the facial nucleus); VLM, ventrolateral reticular nucleus of the medulla; pf-RTN, RTN in a ventromedial position under the facial; cRTN, RTN immediately caudal to the facial nucleus; pFRG, parafacial respiratory group nucleus

**Figure 3.**
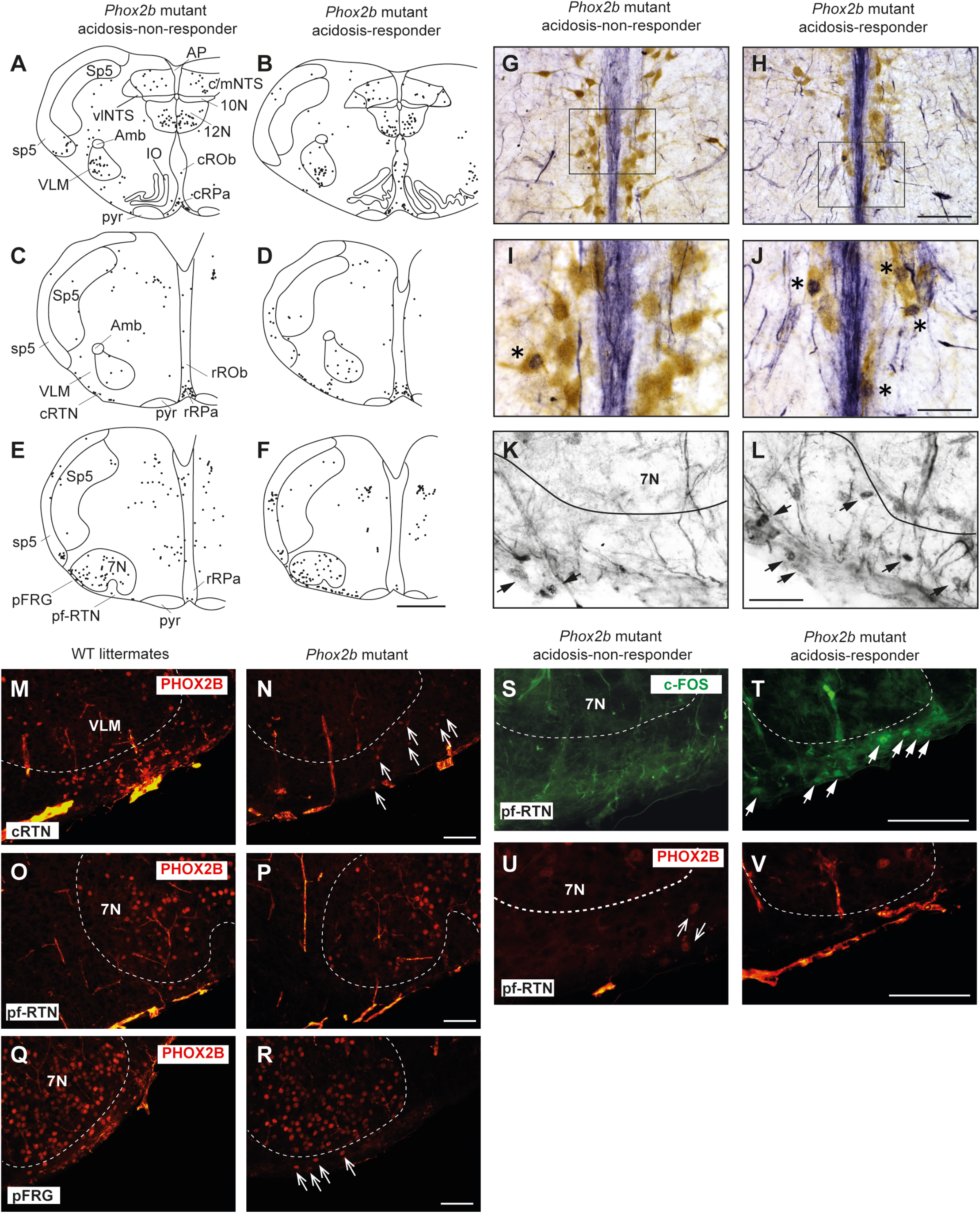
Comparison of the *c-fos* expression between acidosis-responder and acidosis-non-responder *Phox2b* mutant preparations under etonogestrel. Drawings of representative sections from the medulla oblongata at the caudal level (with inferior olives; **A, B**), intermediate level (between inferior olives and the facial nucleus; **C, D**) and rostral level (with the facial nucleus; **E, F**) illustrating the c-FOS distribution in non-acidosis responder (**A, C, E**) and acidosis-responder (**B, D, F**) *Phox2b* mutants under etonogestrel. Photomicrographs of c-FOS/5-HT immunoreactivities in the caudal ROb of acidosis-non responder (**G, I**) and acidosis-responder (**H, J**) *Phox2b* mutants. Photomicrographs in **I** and **J** correspond to enlargements of the outlined area in **G** and **H**; note the greater number of dual-labeled c-FOS/5-HT neurons (asterisks) in acidosis-responder than in non-acidosis responder *Phox2b* mutants. Photomicrographs of c-FOS immunoreactivity in the ventral medullary surface, ventromedial to the facial nucleus in the pf-RTN (**K, L**) showing that the number of c-FOS positive neurons (solid black arrows) was greater in acidosis-responder than in non-acidosis-responder *Phox2b* mutants. Photomicrographs of immunofluorescence detection for PHOX2B in wildtype littermates (**M, O, Q**) and *Phox2b* mutants (**N, P, R**) in the ventral medullary surface, just below the caudal edge of the facial nucleus, in the cRTN (**M, N**), ventromedial to the facial nucleus, in the pf-RTN (**O, P**) and ventrolateral below the facial nucleus, in the pFRG (**Q, R**). In wildtype littermates, PHOX2B neurons are distributed in all three delimitations with the highest number of neurons in the cRTN as already described (Dubreuil et al., 2008). In *Phox2b* mutants, the number of neurons was drastically reduced but some residual cells were still present as previously described (hollow white arrows) (Ramanantsoa et al., 2011). Dual c-FOS (**S, T**) and PHOX2B (**U, V**) detections by immunofluorescence under the ventromedial part of the RTN (pf-RTN) in acidosis-non-responder (**S, U**) and acidosis-responder (**T, V**) *Phox2b* mutants; note the presence of c-FOS positive (solid white arrows) but PHOX2B negative cells in acidosis-responder *Phox2b* mutants. Scale bar = 200 μm (**A, B, C, D, E, F**), 100 µm (**M, N, O, P, Q, R**), 50 μm (**G, H**), and 20 µm (**I, J, K, L, S, T, U, V**). 7N, facial nucleus; 10N, dorsal motor nucleus of the vagus 12N, hypoglossal nucleus; Amb, ambiguus nucleus; AP, area postrema; c/mNTS, commissural and median parts of the nucleus of the tractus solitarius; IO, inferior olives; VLM, ventrolateral medullary reticular nucleus; pFRG, parafacial respiratory group; pyr, pyramidal tract; ROb, raphe obscurus nucleus (caudal part—cROb, from pyramidal decussation to rostral edge of the inferior olive and rostral part—rROb, from rostral edge of the inferior olives to rostral edge of the facial nucleus); RPa, raphe pallidus nucleus (caudal part—cRPa, from pyramidal decussation to rostral edge of the inferior olive and rostral part—rRPa, from rostral edge of the inferior olives to rostral edge of the facial nucleus); RTN, retrotrapezoid nucleus (parafacial RTN—pf-RTN, in ventromedial position under the facial nucleus and caudal RTN—cRTN, immediately to the caudal edge of the facial nucleus); sp5, spinal trigeminal tract; Sp5, spinal trigeminal interpolaris nucleus; vlNTS, ventrolateral part of the nucleus of the tractus solitaries; VLM, ventrolateral medullary reticular nucleus; WT, wildtype.

Under etonogestrel, acidosis-responder *Phox2b* mutants displayed a greater number of c-FOS cells in the *raphe obscurus* nucleus (ROb; **Table 1, Figure 3A, 3B**). This increase concerned more particularly the part of the ROb at the rostro-caudal level delimited by the presence of inferior olives (**Table 1, Figure 3A, 3B**). ROb contains different cell types, including 5-HT neurons (Corcoran et al., 2009, Richerson, 2004); we observed a significant increase in the number of doubly marked c-FOS/5-HT neurons of acidosis-responder *Phox2b* mutants compared to acidosis-non-responders (13.0 [3.5; 29.5] vs 1.0 [0.0; 2.0], p<0.05; **Figure 3G-J**). It should be noted that the difference in c-FOS/5-HT cells in ROb was not due to a different number of 5-HT neurons (788.0 [523.0; 982.0] vs 977.0 [659.0; 1015.0], p<0.05 respectively), a finding that is aligned with the literature (Dubreuil et al., 2008). In the adjacent *raphe pallidus* nucleus, we did not observe a significant difference either in the number of c-FOS (**Table 1**; **Figure 3A-F**) or c-FOS/5-HT neurons (32.5 [13.0; 44.5] vs 17.5 [11.3; 37.0], p>0.05 respectively) between acidosis-responder and acidosis-non-responder *Phox2b* mutants.

In line with published data (Ramanantsoa et al., 2011), we found residual PHOX2B cells in the RTN of *Phox2b* mutants (**Figure 3M-R**). Their number was, expectedly, significantly lower (55.0 [39.0; 98.0]) than in wildtype littermates (291.0 [229.0; 326.0], n=3; p<0.0001). The RTN is described as a short column of relatively sparse neurons in the ventromedial position under the facial nucleus in its rostral part and also immediately to it in its caudal part (Dubreuil et al., 2008, Voituron et al., 2006), named respectively hereafter parafacial RTN (pf-RTN) and caudal RTN (cRTN). In the newborn, RTN is considered to be intertwined with the parafacial respiratory group (pFRG; expiratory generator) that is ventrolaterally below the facial nucleus and also contains PHOX2B cells (Onimaru and Homma, 2003, Onimaru et al., 2008, Onimaru et al., 2014). We thus examined pf-RTN and cRTN as defined above, and pFRG. We did not observe a significant difference in residual PHOX2B cells between acidosis-responder and acidosis-non-responder *Phox2b* mutants in pf-RTN (15 [7; 24] vs 13.5 [5.5; 21.5] respectively), cRTN (27 [17; 51] vs 16.5 [19.5; 33.5] respectively) and pFRG (15.5 [15; 23] vs 12 [5; 16.5] respectively). Yet, we found that acidosis-responder *Phox2b* mutants displayed an increase in the number of c-FOS cells in the pf-RTN (**Table 1, Figure 3E, 3F, 3K, 3L**), but showed no difference at the level of cRTN and pFRG (**Table 1, Figure 3C-F**). Because there was virtually no c-FOS/PHOX2B cells in the pf-RTN, the increase in the number of c-FOS cells in pf-RTN was from PHOX2B-negative cells only (**Figure 3S-2V**).

In other respiratory-related areas of the medulla oblongata, *i*.*e*. the ventrolateral reticular nucleus of the medulla, a part of the reticular formation that contains the ventral respiratory column and the commissural and median part of the nucleus of the solitary tract, we did not observe any difference between acidosis-responder and acidosis-non-responder *Phox2b* mutants (**Table 1, Figure 3A-F**).

### Impact of 5-HT signaling on the respiratory response to metabolic acidosis under etonogestrel

Since 5-HT neurons of the ROb appear to play a key role in restoring the response to acidosis under etonogestrel, we sought to determine whether pharmacological modification of 5-HT signaling modulated the effect of etonogestrel.

#### An increase in 5-HT signaling associated with etonogestrel leads to a potentiation of the respiratory response to acidosis in wildtype Swiss OF1 mice

Preparations from OF1 mice displayed a respiratory-like rhythm in normal-pH of 8.6±0.2 cycles.min^-1^ (n=182) and the classical respiratory response to metabolic acidosis with a respiratory-like rhythm of 14.6±1.0 cycles.min^-1^ (n=14, +40%, p<0.001) as shown in literature (Joubert et al., 2016a, Voituron et al., 2010). As already described in medullary-spinal cord preparations from rodents free from CCHS, etonogestrel (5.10^−2^ µM) increased the respiratory-like rhythm in normal-pH (**Table 2**), but did not modify the increase in fr induced by metabolic acidosis (**Table 2, Figure 1, Figure 4E, 4H**) (Joubert et al., 2016b, Loiseau et al., 2019, Loiseau et al., 2014).

**Table 2.**
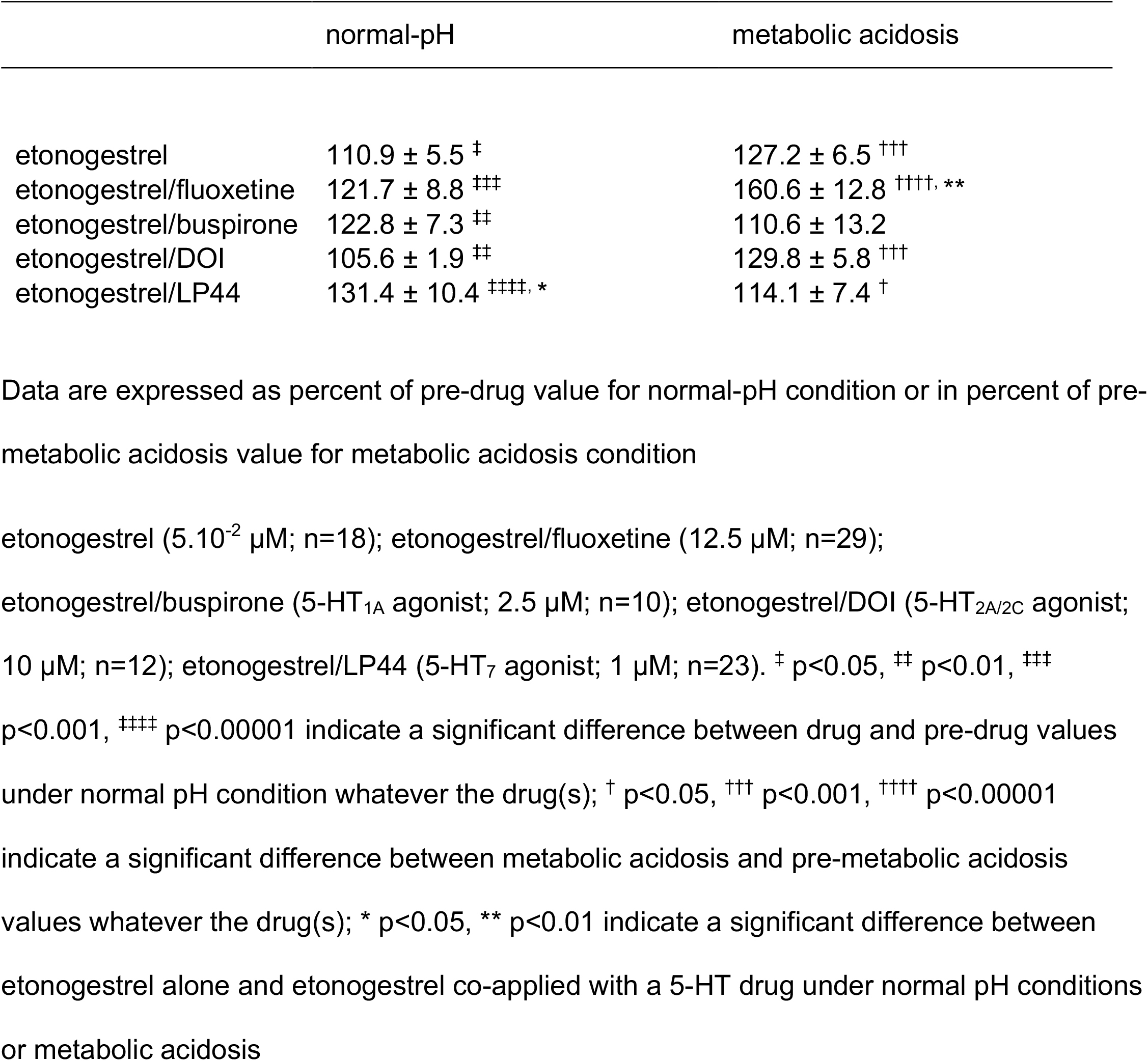
fr in percent of pre-drug or pre-metabolic acidosis in wildtype Swiss OF1 mice

**Figure 4.**
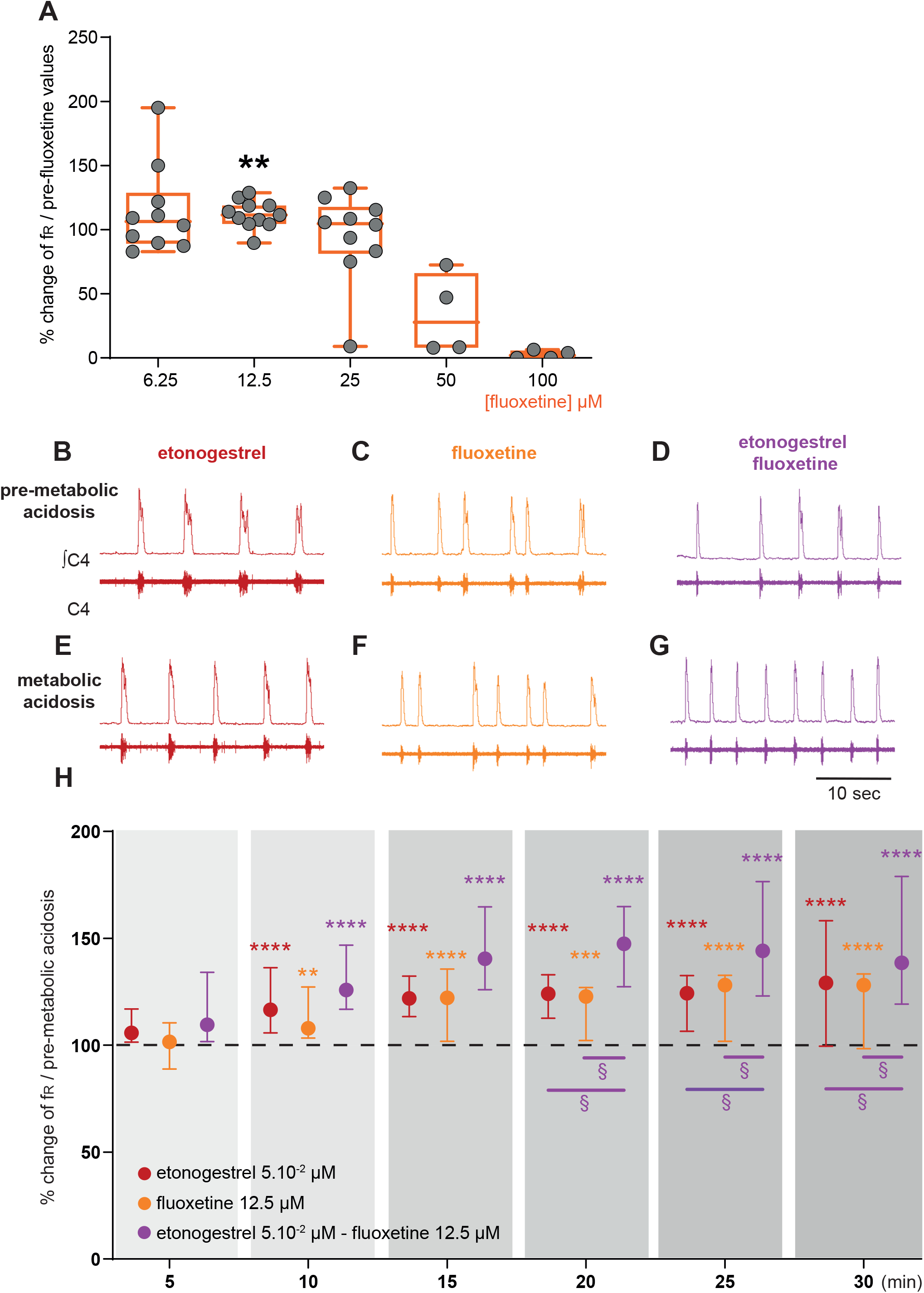
Effect of an increase in 5-HT signaling by fluoxetine on the respiratory response to metabolic acidosis under exposure to etonogestrel in OF1 mice. **A**, Scatter plots with a superimposed box and whisker plot (median [Q1; Q3]) showing percentage of change of fr under 6.25 (n=10), 12.5 (n=11), 25 (n=10), 50 (n=4) and 100 µM (n=4) of fluoxetine. ** p < 0.01 indicates a significant change in fr relative to pre-fluoxetine values (paired t-test or Wilcoxon matched pairs signed rank test depending on the normality of data distribution). **B-G**, Individual phrenic activity traces (fourth cervical ventral nerve root, C4) and integrated C4 activity from medullary-spinal cord from OF1 mice in normal-pH (**B**,**C**,**D**; pre-metabolic acidosis) and metabolic acidosis (**E**,**F**,**G**) conditions, under etonogestrel (**B**,**E**), fluoxetine (**C**,**F**), or etonogestrel/fluoxetine (**D**,**G**) exposure. **H**, Median with interquartile range [Q1; Q3] illustrating the respiratory-like rhythm (respiratory frequency, fr) during the metabolic acidosis challenge per 5-min window under etonogestrel (red circles, n=18), fluoxetine (orange circles, n=11), and etonogestrel/fluoxetine (purple circles, n=29) exposure. Under etonogestrel/fluoxetine, preparations displayed a powerful potentiation of the respiratory response to metabolic acidosis compared to etonogestrel or fluoxetine alone. * Indicates a significant difference in fr relative to pre-metabolic acidosis values (** p < 0.01, *** p < 0.001, **** p < 0.0001; one-way analysis of variance or Friedman test followed by Benjamini, Krieger and Yekutieli’s multiple comparison test). ^§^ Indicates a significant difference (^§^ p < 0.05; two-way analysis of variance test followed by Benjamini, Krieger and Yekutieli’s multiple comparison test) between etonogestrel/fluoxetine preparations and etonogestrel or fluoxetine preparations alone. ∫C4: integrated activity of C4 ventral nerve root; C4: electrical activity of the C4 ventral nerve root.

We first determined the concentration of fluoxetine to be applied as that which does not cause depression of fr. As it has been shown that fluoxetine decreases fr at high concentrations and has no effect at low concentrations in rodent newborns (Bravo et al., 2016, Voituron et al., 2010, Morin et al., 1991), we tested different concentrations: 6.25; 12.5; 25; 50 and 100 µM. At higher concentrations, fluoxetine led to a drop in fr, which in some cases, led to a rhythm arrest (**Figure 4A**). At 25 and 6.25 µM fluoxetine had no significant influence on fr (**Figure 4A**). At 12.5 µM, fluoxetine led to a significant increase in fr (**Figure 4A**). We therefore selected 12.5 µM for rest of the experiments. Then, we observed that the increase in fr induced by metabolic acidosis remained unchanged under fluoxetine (**Figure 1, Figure 4F, 3H, Table 2**).

We then co-applied etonogestrel with fluoxetine (**Figure 1**). In normal-pH, fluoxetine did not change the etonogestrel-induced increase in fr (**Table 2**), and etonogestrel/fluoxetine led to a powerful potentiation of the respiratory response to metabolic acidosis (**Figure 4G, 3H, Table 2**).

We searched for 5-HT receptor(s) involved in the respiratory effect observed under etonogestrel/fluoxetine (**Figure 1**). We selected the 5HT_1A_, 5HT_2A/C_, and 5HT_7_ receptors because they have been identified as the main responsible for the respiratory effects of 5-HT (Hodges and Richerson, 2008), but we did not observe the powerful potentiation of the respiratory response to metabolic acidosis observed with fluoxetine under etonogestrel/5-HT receptor agonists (**Table 2**). Under co-application of etonogestrel with buspirone (5HT_1A_ agonist; 2.5 µM), while OF1 *ex vivo* preparations increase their fr in normal-pH conditions as under etonogestrel alone, they did not show the significant increase in fr characterizing the respiratory response to metabolic acidosis (**Table 2**). Etonogestrel/DOI co-application (5HT_2A/C_ agonist; 10 µM) had same effect as etonogestrel alone with an increase in fr in normal-pH conditions and the classical increase of the respiratory-like rhythm in metabolic acidosis (**Table 2**). Co-application of etonogestrel with LP44 (5HT_7_ agonist; 1 µM) resulted in a greater increase in fr in normal-pH conditions than etonogestrel alone but there was no response to metabolic acidosis in presence of LP44 acidosis (**Table 2**).

#### Increasing the 5-HT signaling in Phox2b mutant mice did not have a positive effect

Considering the data obtained in OF1, we applied fluoxetine at a predefined concentration of 12.5 µM in *Phox2b* mutants and wildtype littermates: it caused a significant decrease in fr in *Phox2b* mutants (0.5±0.1 cycles.min^-1^ vs 1.0±0.2 cycles.min^-1^, -37.7%, p=0.02; **Figure 5A**), an effect observed in OF1 when fluoxetine was applied at higher concentrations, and had no significant effect in wildtype littermates (12.8±2.4 cycles.min^-1^ vs 10.8±1.7 cycles.min^-1^, p=0.25; **Figure 5B**). We decreased the concentration to avoid its depressive effect on the respiratory-like rhythm. At 6.25 and 3.125 µM of fluoxetine, we did not observe a depressive effect in *Phox2b* mutant mice (0.8±0.1 and 0.9±0.2 cycles.min^-1^, in 3.125 and 6.25 µM of fluoxetine respectively *vs* 0.8±0.1 and 0.9±0.2 cycles.min^-1^ without fluoxetine; **Figure 5A**) and their wildtype littermates (13.7±2.2 and 10.6±1.4 cycles.min^-1^, in 3.125 and 6.25 µM of fluoxetine respectively *vs* 13.9±2.5 and 9.4±1.0 cycles.min^-1^ without fluoxetine; **Figure 5B**). We decided to apply fluoxetine at 3.125 µM.

**Figure 5.**
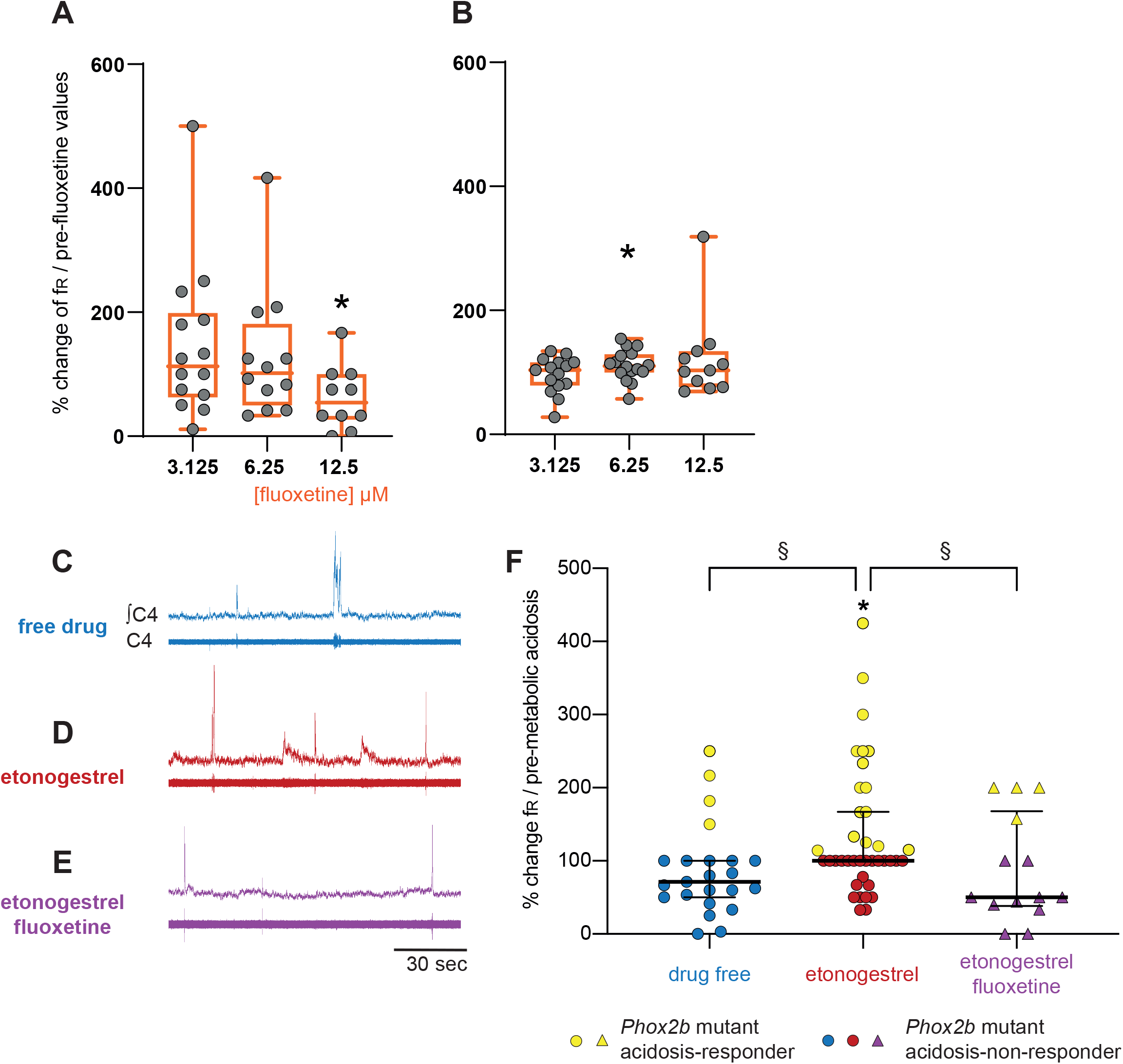
Effect of an increase in 5-HT signaling by fluoxetine in *Phox2b* mutant mice. Scatter plots with a superimposed box and whisker plot (median [Q1; Q3]) showing percentage of change of fr under fluoxetine in *Phox2b* mutants (**A**) and wildtype littermates (**B**). In *Phox2b* mutants, 3.125 (n=14), 6.25 (n=12) and 12.5 µM (n=10) of fluoxetine. In wildtype littermates, 3.125 (n=15), 6.25 (n=16) and 12.5 µM (n=15) of fluoxetine. * p < 0.05 indicates a significant change in fr relative to pre-fluoxetine values (paired t-test or Wilcoxon matched pairs signed rank test depending on the normality of data distribution). **C-E**, Individual phrenic activity traces (fourth cervical ventral nerve root, C4) and integrated C4 activity from medullary-spinal cord from *Phox2b* mutant mice in metabolic acidosis under drug-free (**C**, n=23), etonogestrel (**D**, n=39) and etonogestrel/fluoxetine (**E**, n=14) exposure. **F**, Scatter plot with surperimposed median [Q1; Q3] showing the respiratory-like rhythm (respiratory frequency, fr) during the last five minutes of metabolic acidosis in percentage of pre-metabolic acidosis. In the presence of fluoxetine, the restoration of the respiratory response to metabolic acidosis induced by etonogestrel in *Phox2b* mutant mice was abolished. * Indicates a significant change in fr relative to pre-metabolic acidosis values (* p < 0.05; paired t-test). ^§^ Indicates a significant difference between the 3 conditions (^§^ p < 0.05^;^ Kruskal-Wallis test followed by Benjamini, Krieger and Yekutieli’s multiple comparison test). ∫C4, integrated activity of C4 ventral nerve root; C4, electrical activity of the C4 ventral nerve root. Yellow filled circles or triangles represent *Phox2b* mutant mice acidosis-responders (+10% above pre-metabolic values). Blue filled circles, red filled circles and purple filled triangles represent *Phox2b* mutant mice acidosis-non-responders under, drug-free, etonogestrel and etonogestrel/fluoxetine, respectively.

Under normal-pH, etonogestrel/fluoxetine led to the loss of the effect of etonogestrel on fr in wildtype littermates (14.0 ± 2.8 vs 13.7 ± 2.2 cycles.min^-1^ free of drugs, p=0.43) and had no effect in *Phox2b* mutants (0.7 ± 0.1 vs 0.8 ± 0.1 cycles.min^-1^, p=0.42). Under metabolic acidosis, opposite to what we expected, the co-application of drugs did not result in potentiation of the respiratory response to metabolic acidosis in wildtype littermates (+45% *vs* +28%, p=0.46) and the restoration of this response observed under etonogestrel alone in *Phox2b* mutants was not present (**Figure 5E, 4F**).

### Characterization of 5-HT metabolic pathways in *Phox2b* mutants, wildtype littermates and OF1 mice

We compared the 5-HT metabolic pathways of the medulla oblongata between OF1, *Phox2b* mutants and wildtype littermates to investigate whether differences could contribute to the observed discrepancy in the respiratory effects of etonogestrel. While medullary 5-HT quantity was lower in *Phox2b* mutants and wildtype littermates than in OF1 (**Figure 6B**), quantities of 5-HTP (5-HT precursor) and 5-HIAA (5-HT degradation product) were higher in wildtype littermates than in OF1, and were similar between *Phox2b* mutants and OF1 mice. The 5-HT/5-HTP ratio was lower in *Phox2b* mutants and wildtype littermates than in OF1 (**Figure 6C**), indicating a weak 5-HT synthesis in *Phox2b* mutants and wildtype littermates. The ratio 5-HIAA/5-HT was higher in *Phox2b* mutants and wildtype littermates than in OF1 (**Figure 6D**), indicating a higher 5-HT turn-over.

**Figure 6.**
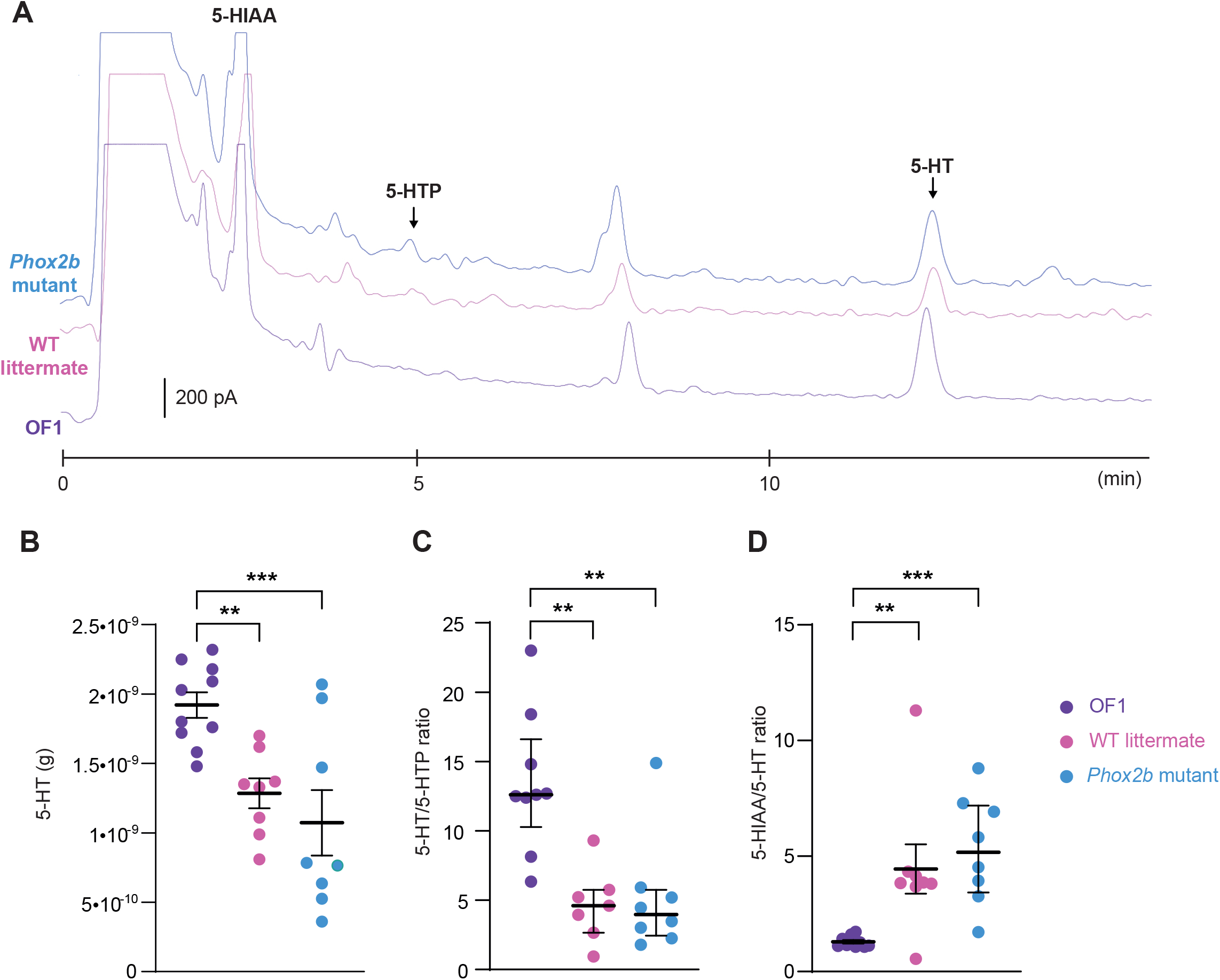
Ultra-high-performance liquid chromatography analysis of 5-HT concentration and its related compounds, 5-HTP and 5-HIAA, in the medulla oblongata of *Phox2b* mutants, wildtype littermates and OF1 mice. **A**, Example of a chromatogram of medulla oblongata obtained from *Phox2b* mutants (blue; n=8), wildtype littermates (pink; n=8) and OF1 mice (purple; n=10) with indicated peaks of serotonin (5-HT), its precursor 5-hydroxytryptophan (5-HTP) and its metabolite 5-hydroxyindole acetic acid (5-HIAA). **B**, Scatter plots showing 5-HT quantity (g of 5-HT per medulla) with superimposed mean ± standard error of the mean. **C** and **D**, Scatter plots showing, respectively, 5-HT/5-HTP and 5-HIAA/5-HT ratio with median [Q1; Q3] superimposed in OF1 (purple filled circles), wildtype littermates (pink filled circles) and *Phox2b* mutants (blue filled circles). * Indicates a significant difference between OF1, *Phox2b* mutants and wildtype littermates (** p < 0.01 and *** p < 0.001; one-way analysis of variance or Kruskal-Wallis test followed by Benjamini, Krieger and Yekutieli’s multiple comparison test).

## DISCUSSION

Here, in a preclinical model of CCHS, we showed that etonogestrel induces a restoration of the central H^+^ chemosensitivity in some preparations. The etonogestrel-induced effects seem to depend on the functional status of medullary serotoninergic systems but not on residual PHOX2B cells in RTN, where structural alterations are considered to be the main cause of central hypoventilation attributed to reduction or loss of CO_2_/H^+^ sensitivity in CCHS patients (Ramanantsoa et al., 2011, Dubreuil et al., 2008).

### Etonogestrel-induced restoration of central H^+^ chemosensitivity, a respiratory effect not systematically present in *Phox2b* mutant mice

The present study is the first to report a restoration of H^+^ chemosensitivity in a preclinical model of CCHS, confirming previous clinical observations with desogestrel (Straus et al., 2010). Putting these observations into perspective with the small amount of data available in the literature suggests that etonogestrel either has a specific action compared to progesterone or other progestins, or has enhanced activity compared to progesterone or medroxyprogesterone. As noted the ventilatory response to CO2 of CCHS patients is not increased during pregnancy (Sritippayawan et al., 2002) and medroxyprogesterone, a pregnane progestin, does not improve ventilation in CCHS patients (Weese-Mayer et al., 1992).

We observed marked heterogeneity in the response of *Phox2b* mutants to metabolic acidosis, leading to separate the preparations in acidosis-responders and acidosis-non-responders. Few *Phox2b* mutants were categorized as acidosis-responders in the absence of progestin, consistent with the notion that the absence or dysfunction of PHOX2B RTN cells abolishes or nearly abolishes CO_2_/H^+^ chemosensitivity (Ramanantsoa et al., 2011, Dubreuil et al., 2008, Guyenet and Bayliss, 2015, Gestreau et al., 2010). Yet, the very existence of acidosis-responder preparations suggests the possible persistence of H^+^ chemosensitive mechanisms despite *Phox2b* mutations in the RTN. Such a phenomenon could contribute to the residual chemosensitivity described in some CCHS patients (Carroll et al., 2014). Etonogestrel could thus activate or over-activate these residual mechanisms, which would be in line with our observations of both a consistent restoration of the response to metabolic acidosis across all preparations and of an increase in the proportion of acidosis-responders. Whatever the mechanisms involved (etonogestrel-related activation or over-activation of residual RTN sensitivity, or activation of pathways that are silent without progestin), our data show that the *ex vivo* ventilatory-like effects of etonogestrel are not systematically observed, in a manner similar to clinical effects in CCHS patients (Straus et al., 2010, Li, 2013). This suggests that the effects of etonogestrel depend on cell or molecular targets whose functional status is variable among individuals.

### 5-HT neurons, but not PHOX2B residual RTN cells, as a neural basis for the recovery of the CO_2_/H^+^ chemosensitivity in *Phox2b* mutants

Under etonogestrel, the greater number of ROb 5-HT neurons expressing *c-Fos* in acidosis-responder *Phox2b* mutants compared to acidosis-non-responders led us to suppose that 5-HT neurons play a key role. This agrees with previous observations in CCHS-free rodents (Joubert et al., 2016b, Loiseau et al., 2018). Many *ex vivo* and *in vivo* studies support the hypothesis that 5-HT neurons within the medullary raphe act as CO_2_/H^+^ chemoreceptors and contribute a ventilatory response appropriate to maintain homeostasis (Corcoran et al., 2009, Richerson, 2004, Bernard et al., 1996, Depuy et al., 2011, Peever et al., 2001, Holtman et al., 1986). In particular, the activity of ROb 5-HT neurons that heavily innervate respiratory-related structures is increased under hypercapnia, increasing breathing frequency, and potentiating the CO_2_/H^+^ respiratory chemoreflex (Depuy et al., 2011, Holtman et al., 1986, Veasey et al., 1995). We therefore believe that, in response to etonogestrel, ROb 5-HT neurons contribute to the restoration of an acidosis response in *Phox2b* mutants through a release of 5-HT within respiratory structures. In this hypothesis, these ROb 5-HT neurons would be more sensitive to an etonogestrel effect in the acidosis-responder individuals. The action of etonogestrel on ROb 5-HT neurons could proceed from their depolarization independently of metabolic acidosis, in turn leading to a greater response to the acid challenge. Alternatively, etonogestrel could directly increase the sensitivity of these neurons to acidosis. In either case, the molecular target of etonogestrel remains to be identified.

Another possible explanation for the restoration of the acidosis-induced increase in respiratory drive could be the activation of RTN residual PHOX2B cells. Etonogestrel modulates the expression of *PHOX2B* and its target genes, suggesting that it can counteract the loss of function and the toxic effects due to the mutation (Cardani et al., 2018, Di Lascio et al., 2020). However, acidosis-responder and acidosis-non-responder preparations did not differ regarding *c-Fos* expression in RTN PHOX2B cells. This divergence from previous data showing beneficial effects of etonogestrel on RTN PHOX2B cells, which have been observed on neuroblastoma cell lines in the context of prolonged exposure (Cardani et al., 2018, Di Lascio et al., 2020), could stem from differences in experimental conditions. In the present work, the etonogestrel exposure was brief and possibly insufficient to observe genomic effects. Also, neuroblastoma cell lines express nuclear progesterone receptors, which has not been demonstrated in RTN PHOX2B cells even though hypercapnia-induced RTN *c-Fos* expression is higher in females than males (Niblock et al., 2012). Of note, the fact that residual PHOX2B cells did not seem to be involved in restoring a respiratory response to metabolic acidosis in our setting does not mean that this would not be the case in a more integrated organism and/or under longer exposure to etonogestrel. This pathway would however not relate to the effects mediated by the activation of medullary 5-HT neurons, but possibly be additive in nature.

### Importance of 5-HT metabolic pathways in the potentiation of the respiratory response to metabolic acidosis induced by co-application of fluoxetine and etonogestrel

An interaction between etonogestrel and 5-HT systems is supported by the fact that potentiation of the response to metabolic acidosis under etonogestrel was only observed, in OF1 mice, when 5-HT systems were boosted by fluoxetine. It can therefore be assumed that the effects of etonogestrel involves excitation of 5-HT neurons leading to 5-HT release, in turn responsible for the enhancement of the respiratory drive. In OF1 mice whose RTN PHOX2B neurons are functional, this hypothesis implies that the etonogestrel-induced 5-HT release would not modulate the respiratory response to metabolic acidosis, which is consistent with published data at this stage of development (Corcoran et al., 2009, Hodges and Richerson, 2008). In contrast, etonogestrel combined with a fluoxetine-induced excess of 5-HT in the synaptic cleft would lead to a reinforcement of the respiratory response to metabolic acidosis, which suggests the need for a large quantity of 5-HT to be released for the respiratory effects of etonogestrel to become visible. In this case, it could be assumed that 5-HT was released in large quantities in *Phox2b* mutant mice to exert a stimulating effect. This would allow for the compensation of the reduced number of RTN PHOX2B cells and/or the mutation-related dysfunction of residual RTN PHOX2B cells. Our ultra-high-performance liquid chromatography data, showing lower 5-HT levels in *Phox2b* mutants than in OF1 individuals, make a larger release of 5-HT in *Phox2b* mutants unlikely. It is possible, as already reported in another context (Husch et al., 2012), that *Phox2b* mutants have a supersensitivity to 5-HT due to an increased number/functionality of 5-HT receptors. This would allow the etonogestrel-induced 5-HT release to exert a greater stimulating effect on the respiratory network, hence an enhanced fr in metabolic acidosis. It should be noted that since the low quantity of 5-HT was also found in wildtype littermates, the difference between *Phox2b* mutants and the classically used wildtype Swiss OF1 strain did not depend on the mutation but on the strain, as already described (Menuet et al., 2011).

The high potency of 5-HT systems in *Phox2b* mutants, irrespective of its origin, may explain the apparently paradoxical detrimental effect of the co-application of fluoxetine with etonogestrel. Indeed, 5-HT can exert a depressant effect on the respiratory drive at high concentrations (Bravo et al., 2016, Voituron et al., 2010, Morin et al., 1991). Under etonogestrel/fluoxetine co-application, the quantity of 5-HT within the respiratory network could be too high, leading to a depressant effect that cancelled the stimulating effect of etonogestrel. This highlights the importance of the functional status of 5-HT systems for an etonogestrel-induced respiratory effect to occur in a CCHS context. This could explain why some CCHS patients are sensitive to etonogestrel regarding the ventilatory response to CO_2_ while others are not (Straus et al., 2010, Li, 2013). This hypothesis is all the more worth testing given that the known polymorphisms related with serotonergic neurotransmission (Heils et al., 1996) have been involved in the clinical expression of diseases (Eddahibi et al., 2003), and the reactions to certain treatments (Bocchio-Chiavetto et al., 2008). Under this hypothesis, co-administration of etonogestrel with fluoxetine or another selective serotonin reuptake inhibitor could be considered if etonogestrel alone does not produce ventilatory effects, and if 5-HT systems can be characterized as less potent in etonogestrel-insensitive patients compared to etonogestrel-sensitive ones.

To conclude, regardless of the precise molecular pathways involved, collective data point to the potential for targeting 5-HT signaling as a therapeutic intervention associated with etonogestrel for improving the respiratory drive of CCHS patients. The fact that etonogestrel is approved for clinical use for contraception and fluoxetine or other selective serotonin reuptake inhibitors are approved as anti-depressive drugs lends support to translational opportunities.

## METHODS

### Ethical approval

Experiments were carried out in accordance with Directive 2010/63/EU of the European Parliament and of the Council of 22 September 2010 French law (2013/118). Protocols were approved by Charles Darwin Ethics Committee for Animal Experimentation (Ce5/2011/05; APAFIS#14259-2018032518034654v3 and #2210-2015100812195835v2) and all efforts were made to minimize the number of animals used and their suffering.

### Animals

Experiments were performed on newborn mice (0-4 days old): *Egr2*^*cre/+*^; *Phox2b*^*27Ala/+*^ mutant mice (hereafter termed *Phox2b* mutants; 1.2±0.02 g, n=106), wildtype littermates (1.2±0.02 g, n=123) and wildtype Swiss OF1 strain (2.3±0.05 g, n=192; Charles River laboratories, L’Arbresle, France).

Mutant mice expressing a Congenital Central Hypoventilation Syndrome-causing expansion of the 20-residue poly-alanine stretch in *Phox2b* were generated by crossing *Egr2*^*cre/+*^ males with *Phox2b*^*20-27Ala/20-27Ala*^ females (Voiculescu et al., 2000, Ramanantsoa et al., 2011). Genotyping was done on tail DNA to identify mutants and wildtype littermates. To detect *cre* for mutant mice identification, the primers AAATTTGCCTGCATTACCG and ATGTTTAGCTGG CCCAAATG were used, yielding a band of 250 pb.

### Drugs

Etonogestrel (3-ketodesogestrel), fluoxetine, buspirone, DOI (1-(2,5-dimethoxy-4-iodophenyl)-2-aminopropane hydrochloride-hydrochloride) and serotonin were purchased from Sigma-Aldrich (Saint-Quentin Fallavier, France). LP44 (4-[2-(methylthio)phenyl]-N-(1,2,3,4-tetrahydro-1-naphthalenyl)-1-piperazinehexanamide hydrochloride) was purchased from Tocris Bioscience (Bristol, UK). Drugs were prepared in saline except for etonogestrel that was dissolved in dimethylsulfoxide. All drugs were dissolved in artificial cerebrospinal fluid with a final concentration dimethylsulfoxide at 0.01% if appropriate.

### *Pharmacology on ex vivo* medullary-spinal cord preparation

#### Medullary-spinal cord preparations

Newborn mice were placed under deep cold anesthesia (Danneman and Mandrell, 1997) and medullary-spinal cord preparations were dissected out, as previously described (Suzue, 1984, Joubert et al., 2016b, Voituron et al., 2011). The rostral section was made at the level of the anterior inferior cerebellar arteries just caudal to the eighth cranial nerve exit points. The caudal section was made between the seventh and the eighth cervical spinal roots. *Ex vivo* preparations were placed in a recording chamber with the ventral surface facing upward. They were superfused with artificial cerebrospinal fluid (in mM: 129 mM NaCl, 3.35 mM KCl, 1.26 CaCl_2_, 1.15 mM MgCl_2_, 0.58 mM NaH_2_PO_4_, 30 mM Glucose and NaHCO_3_ at various concentrations depending on the experimental condition (Murakoshi et al., 1985) maintained at 27±1°C, saturated with O_2_, and adjusted to the appropriate pH by bubbling with 95% O_2_ and 5% CO_2_. As molecular sensors detecting H^+^ and CO_2_ changes are described as sensitive to an increased concentration of H^+^, we performed pH variation of artificial cerebrospinal fluid to mimic physiological consequences of an increase in CO_2_ (Gestreau et al., 2010, Song et al., 2012, Guyenet and Bayliss, 2015) (Kumar et al., 2015). Normal-pH artificial cerebrospinal fluid (pH 7.4) and metabolic acidosis artificial cerebrospinal fluid (pH7.23) differed in terms of NaHCO_3_ concentration (21 mM and 15 mM, respectively) (Murakoshi et al., 1985, Suzue, 1984, Gestreau et al., 2010, Loiseau et al., 2019). Electrical activity of the fourth cervical ventral nerve root (C4) was recorded using a suction electrode, filtered (300-1000 Hz), amplified (x10000 Grass P511 AC Amplifier), integrated (Dual channel integrator, University of Chicago), and digitized through a Spike 2 data analysis system (CED micro 1401; Cambridge Electronic Design) with a sampling frequency of 2500 Hz. Respiratory-like rhythm (respiratory frequency, fr) was defined as the burst frequency recorded on the fourth cervical ventral nerve root for 1 minute (burst.min^-1^) (Murakoshi et al., 1985, Suzue, 1984, Gestreau et al., 2010, Loiseau et al., 2019).

After surgery, preparations were routinely allowed to stabilize for 30 minutes in normal-pH free of drug (**Figure 1A**). Consistent with previous reports about the respiratory effect of etonogestrel or other steroids and 5-HT drugs on newborn rodents less than 4 days old, we pooled data obtained from males and females (Loiseau et al., 2019, Joubert et al., 2016b, Morin et al., 1990, Voituron et al., 2010, Ren and Greer, 2006). Baseline values were defined as the mean value during the last 5 minutes of this stabilization period. Then, a given preparation was exposed to a given pharmacological protocol consisting of 20 minutes of drug under normal-pH conditions, followed by exposure to metabolic acidosis under the same drug for 30 minutes, followed by a 30-minute washout period. To evaluate the influence of a drug under normal-pH conditions, the last five minutes of drug exposure was compared to the five minutes preceding the drug exposure (**Figure 1A**). The effect of a drug on the respiratory metabolic acidosis response was assessed by comparing the last five minutes before metabolic acidosis (pre-metabolic value) to the last five minutes of the metabolic acidosis (**Figure 1A**). To compare between experimental conditions (animals and drugs), we normalized fr values during metabolic acidosis to the value of the last 5 minutes before metabolic acidosis.

#### Pharmacological exposure to etonogestrel on Phox2b mutants and wildtype littermates

*Phox2b* mutant (n=23) and wildtype littermate (n=31) preparations were exposed only to the drug-free metabolic acidosis protocol to assess whether they showed a respiratory response to metabolic acidosis (**Figure 1B**).

Based on previous work (Joubert et al., 2016b, Loiseau et al., 2019), to determine the respiratory influence of etonogestrel in normal-pH conditions and the respiratory response to metabolic acidosis, 39 preparations of *Phox2b* mutants and 38 preparations of wildtype littermates were exposed to 5.10^−2^ µM of etonogestrel (**Figure 1B**).

#### Pharmacological exposure to etonogestrel co-applied with serotonergic drugs in OF1 mice

We investigated whether the combination of etonogestrel with a selective serotonin reuptake inhibitor, fluoxetine, induced a respiratory benefit. As the respiratory influence of fluoxetine has been shown to be variable according to its concentration in rodent newborns (Morin et al., 1991, Voituron et al., 2010, Bravo et al., 2016), we tested different concentrations to determine a dose that did not cause respiratory depression in OF1 mice. Preparations were superfused for 20 minutes at normal-pH with fluoxetine at 6.25 µM (n =10), 12,5 µM (n=11), 25 µM (n=10), 50 µM (n=4) and 100 µM (n=4); 12.5 µM was retained (see results). Second, preparations were exposed to either 5.10^−2^ µM etonogestrel alone (n=18), 12.5 µM fluoxetine alone (n=11) or both 5.10^−2^ µM etonogestrel and 12.5 µM fluoxetine (n=29) for 20 minutes at normal-pH, followed by 30 minutes under metabolic acidosis with the considered drug and finally 30 minutes under normal-pH free of drug (**Figure 1A, 1B**).

As we observed a benefit from the combination of etonogestrel and fluoxetine, we investigated the 5-HT receptors involved by co-applying etonogestrel with a 5-HT_1A_ agonist (buspirone), a 5-HT_2A/2C_ agonist (DIO), or a 5-HT_7_ agonist (LP44) (Hawkins et al., 2015, Morin et al., 1991, Di Pasquale et al., 1994). First, optimal concentrations were determined as the minimum concentrations inducing a stimulatory effect at normal-pH by examining exposure to the following: buspirone (0.5 [n=9], 2.5 [n=10] and 5 µM [n=6]), DOI (5 [n=11], 10 [n=12] and 20 µM [n=11]) and LP44 (1 µM [n=23]). The optimal concentrations determined were 2.5 µM buspirone,10 µM DOI, and 1 µM LP44. Second, preparations were co-exposed to 5.10^−2^ µM etonogestrel and 5-HT receptor agonists (2.5 µM buspirone [n=10], 10 µM DOI [n=12] and 1 µM LP44 [n=23]) for 20 minutes under normal pH, then 30 minutes under metabolic acidosis, followed by a return under normal-pH free of drugs (**Figure 1A, 1B**).

#### Pharmacological exposure to etonogestrel co-applied with fluoxetine in Phox2b mutants and wildtype littermates

Given the results obtained in OF1 mice preparations, we investigated a respiratory benefit to the combination of etonogestrel with fluoxetine. First, preparations from 10 *Phox2b* mutants and 15 wildtype littermates were exposed to 12.5 µM of fluoxetine at normal-pH. At this concentration, fluoxetine decreased the fr of *Phox2b* mutant mice. We therefore decided to test 3.125 µM (n=14 *Phox2b* and n=15 wildtype littermates) and 6.25 µM (n =12 *Phox2b* and n=16 wildtype littermates) of fluoxetine. 3.125 µM fluoxetine was retained (see results). Second, *Phox2b* mutant (n=14) and wildtype littermates (n=15) preparations were co-exposed to 5.10^−2^ µM etonogestrel and 3.125µM fluoxetine under normal-pH and metabolic acidosis (**Figure 1A, E1B**), followed by a washout with normal-pH free of drug.

### Histology

Immunohistological investigations were performed on acidosis-responder and acidosis-non-responder *Phox2b* mutants exposed to 5.10^−2^ µM etonogestrel as described above with the aim of revealing the cell populations responsible for the different respiratory behavior between the two types of mutants. After the 30-minute period of metabolic acidosis exposure under etonogestrel, preparations were fixed by immersion in 4% paraformaldehyde in 0.1 M phosphate buffer (pH 7.4) for 72 h at 4°C. Then, they were cryoprotected for 48 hours in 0.1 M phosphate buffer containing 30% sucrose and stored at -20°C until use. Free-floating coronal sections (30 µm) obtained using a cryostat (Leica CM 1510S) were used for immunohistological investigations.

Dual immunohistochemical detections of c-FOS and 5-HT were processed. First, sections were incubated with a rabbit polyclonal antibody against c-FOS (sc-253; Santa Cruz Biotechnology Inc., CA, USA, 1:8000, in 1% bovine serum albumin, BSA; 48 h, 4°C), then with a biotinylated goat anti-rabbit IgG antibody (BA-1000, Vector Laboratories, Burlington, Canada; 1:500; in 1 % BSA; 2 h, room temperature), and finally an avidin-biotin-peroxidase complex (ABC kit standard, PK-6100, Vectastain, Vector Laboratories, Burlingame, CA, USA; 1:250; 1 h). Peroxidase activity was detected using a solution containing 0.02 % 3,3’-diaminobenzidine tetrahydrochloride, 0.04 % nickel ammonium sulfate and 0.01 % H_2_O_2_ in 0.05 m Tris-HCl buffer (pH 7.6), which results in a blue/grey chromogen. Second, sections were incubated with a rabbit polyclonal antibody against 5-HT (S8305, Sigma-Aldrich, Saint-Quentin Fallavier, France; 1:20000, in 1 % BSA; 48 h, 4°C). Sections were then incubated with biotinylated goat anti-rabbit (BA-1000, Vector Laboratories, Burlington, Canada; 1:500 in 1 % BSA; 2 h, room temperature), and then ABC (1:250; 1 h). Peroxidase activity was detected using a solution containing 0.02 % 3,3’-diaminobenzidine tetrahydrochloride and 0.01 % H_2_O_2_ in 0.05 m Tris-HCl buffer (pH 7.6), which results in a brown chromogen. Some sections were processed in parallel, but with the omission of primary or secondary antibodies. No labelling was observed in these conditions. All sections were mounted in sequential caudo-rostral order on silanized slides, air-dried, and coverslipped with EUKITT (Bio Optica, Milan, Italy). c-FOS, 5-HT, and c-FOS/5-HT immunolabeled cells were visually counted by an investigator blinded to samples using a light microscope (Leica DM2000; Leica Microsystems, Heidelberg, Germany) at high magnification (x200 or x400 depending on the immunolabeled density of cells). Analysed medullary structures involved in elaboration or adaptation of the respiratory drive were localized using standard landmarks (Paxinos and Franklin, 2001, Paxinos et al., 2007). c-FOS positive cells were counted in commissural and medial parts of the nucleus of the solitary tract, in the ventrolateral medullary reticular nucleus, a neuronal column ventral to nucleus ambiguous, extending from pyramidal decussation to caudal edge of facial nucleus, that contains the ventral respiratory column in medullary raphe nuclei *i*.*e*. the raphe *obscurus*, the raphe *pallidus*, in the ventral medullary surface at the level of the region containing the retrotrapezoid nucleus and the parafacial respiratory group, *i.e*.in ventromedial position under the facial nucleus in the rostral part of the retrotrapezoid nucleus and also immediately to it in its caudal part (Dubreuil et al., 2008, Voituron et al., 2006), named respectively hereafter parafacial retrotrapezoid nucleus and caudal retrotrapezoid nucleus, and ventrolaterally below the facial nucleus and also contains in parafacial respiratory group. Note that a distinction was made between the caudal part of the raphe *pallidus* and raphe *obscurus* (from the pyramidal decussation to the rostral edge of the inferior olive) and their rostral part (from the rostral edge of the inferior olives to the rostral edge of the facial nucleus) (Joubert et al., 2016b). Sections were photographed with a digital camera (Leica DFC450C, Leica Microsystems, Heidelberg, Germany) using the software Leica Application Suite (L.A.S V4.5). c-FOS and c-FOS/5-HT cell counts were performed by eye using a counting gird in the eyepiece of the microscope at x400 to count all immunolabeled cells by varying the micrometer of the microscope, which was essential for tissue sections of 30 µm. We compared the total number of cells for each analysed area between acidosis-responder and acidosis non-responder *Phox2b* mutant preparations.

Dual detections of c-FOS and PHOX2B by immunohistofluorescence were performed. Sections were co-incubated with a rabbit polyclonal antibody against c-FOS (sc-253; Santa Cruz Biotechnology Inc., CA, USA, 1:8000, in 1%, BSA; 48 h, 4°C) and a mouse monoclonal antibody against PHOX2B (sc-376993; Santa Cruz Biotechnology Inc., CA, USA, 1:1000, in 1%, BSA; 48 h, 4°C). Sections were then incubated with an Alexa Fluor 488-labeled donkey anti-rabbit antibody (Molecular Probes, Eugene, OR) and an Alexa Fluor 555-labeled goat anti-mouse antibody (Invitrogen, Thermo Fisher Scientific) concomitantly with DAPI at 1:1000 (Immunochemistry technologies, 2 hours, room temperature). They were then washed, mounted in sequential caudo-rostral order on silanized slides, air-dried, and coverslipped using Fluoromount G (Fluoromount Aqueous Mounting Medium, Sigma-Aldrich, Saint-Quentin Fallavier, France). Sections were examined using a fluorescence microscope (Leica DFC450C, Leica Microsystems, Heidelberg, Germany). c-FOS, PHOX2B and c-FOS/PHOX2B cells were counted in the caudal retrotrapezoid nucleus, parafacial retrotrapezoid nucleus, and parafacial respiratory group. Cell counts were performed on images obtained with a digital camera (Leica DFC450C, Leica Microsystems, Heidelberg, Germany). We compared the total number of cells for each analyzed area between acidosis-responder and acidosis non-responder *Phox2b* mutant preparations.

### Ultra high-performance liquid chromatography neurotransmitter analyses

The levels of 5-HT, its precursor 5-hydroxytryptophan (5-HTP) and its metabolite 5-hydroxyindole acetic acid (5-HIAA) were measured by ultra high-performance liquid chromatography in the medulla oblongata of *Phox2b* mutants (n=8), wildtype littermates (n=8) and OF1 mice (n=10). After rapid cold surgery, the medulla oblongata was taken and kept at - 80°C until analyses (Ramirez and Viemari, 2005). Before ultra high-performance liquid chromatography, tissues were ground by sonication and centrifuged, the supernatant was used to quantify 5-HT and its related compounds, 5-HTP and 5-HIAA. Tissue analyte levels were quantified by ultra high-performance liquid chromatography coupled with electrochemical detection. The ultra high-performance liquid chromatography system consisted of a degasser (Prominence), a high-pressure (LC-30 AD pump) and an autosampler (SIL-30AC Shimadzu). Separations were performed using a 100 × 2.1 mm Kinetex C18 core-shell 2.6 µm column (00D-4462-AN, Phenomenex) equipped with an Ultra Security Guard (AJ0-8782, Phenomenex) as a precolumn. The mobile phase (70 mmol/L potassium phosphate containing 0.1 mmol/L EDTA, 6.6 mmol/L octane sulfonate, 3.1 mmol/L triethylamine, 10% methanol, pH adjusted to 3.12 with 1 mmol/L citric acid, filtered), was pumped at a flow rate of 0.4 mL/min. Analytes were detected at an oxidation potential of 800 mV versus the reference electrode. Chromatograms were acquired at a rate of 10 Hz using Lab Solutions software for 17 minutes per sample. All the samples had to be diluted 3-fold in the extraction solution to properly quantify 5-HIAA. Concentrations of analytes were determined by comparison of chromatographic peak areas with calibration curves derived from a mixture of synthetic standards. Final concentrations were expressed as g of analytes per medulla oblongata and 5-HIAA/5-HT and 5-HT/5-HTP ratios were calculated.

### Statistics

Data were analyzed with GraphPad (GraphPad Prism9, San Diego, California, United-States). Normality of data distribution for fr, for cells immunolabelled c-FOS, 5-HT, c-FOS/5-HT, PHOX2B, c-FOS/PHOX2B, for concentrations of 5-HT, 5-HTP and 5-HIAA, and for 5-HT/5-HTP and 5-HIAA/5-HT ratios were assessed using the d’Agostino and Pearson omnibus normality test. Depending on whether the distribution was normal or not, data were expressed as mean ± standard error of the mean or median and interquartile range [Q1; Q3] and parametric or non-parametric tests were performed: within a group of preparation, in paired conditions, paired t-test, Wilcoxon signed rank test, one-way analysis of variance, or Friedman test followed by Dunn’s or Benjamini, Krieger and Yekutieli’s multiple comparison test; between two groups, in unpaired conditions, unpaired t test, Mann & Whitney test and two-way analysis of variance followed by Benjamini, Krieger and Yekutieli’s multiple comparison test in unpaired conditions; between more than two groups, one way analysis of variance or Friedman test followed by Benjamini, Krieger and Yekutieli’s multiple comparison test. Percentages of *Phox2b* mutant mice acidosis-responder or acidosis-non-responder with or without etonogestrel were compared by Fisher’s exact test. Differences were considered to significant if p < 0.05.

## ACKOWLEDGEMENTS

We thank Andrew Lane PhD (Lane Medical Writing) for editing the English text.

## REFERENCES

Amiel, J., Dubreuil, V., Ramanantsoa, N., Fortin, G., Gallego, J., Brunet, J. F. & Goridis, C. 2009. PHOX2B in respiratory control: lessons from congenital central hypoventilation syndrome and its mouse models. Respir Physiol Neurobiol., 168, 125–132.

Amiel, J., Laudier, B., Ttie-Bitach, T., Trang, H., De, P. L., Gener, B., Trochet, D., Etchevers, H., Ray, P., Simonneau, M., Vekemans, M., Munnich, A., Gaultier, C. & Lyonnet, S. 2003. Polyalanine expansion and frameshift mutations of the paired-like homeobox gene PHOX2B in congenital central hypoventilation syndrome. Nat Genet., 33, 459–461.

Bernard, D. G., Li, A. & Nattie, E. E. 1996. Evidence for central chemoreception in the midline raphe. J Appl Physiol (1985), 80, 108–15.

Bocchio-Chiavetto, L., Miniussi, C., Zanardini, R., Gazzoli, A., Bignotti, S., Specchia, C. & Gennarelli, M. 2008. 5-HTTLPR and BDNF Val66Met polymorphisms and response to rTMS treatment in drug resistant depression. Neurosci Lett, 437, 130–4.

Bravo, K., Eugenin, J. L. & Llona, I. 2016. Perinatal Fluoxetine Exposure Impairs the CO2 Chemoreflex. Implications for Sudden Infant Death Syndrome. Am J Respir Cell Mol Biol, 55, 368–76.

Cardani, S., Di Lascio, S., Belperio, D., Di Biase, E., Ceccherini, I., Benfante, R. & Fornasari, D. 2018. Desogestrel down-regulates PHOX2B and its target genes in progesterone responsive neuroblastoma cells. Exp Cell Res, 370, 671–679.

Carroll, M. S., Patwari, P. P., Kenny, A. S., Brogadir, C. D., Stewart, T. M. & Weese-Mayer, D. E. 2014. Residual chemosensitivity to ventilatory challenges in genotyped congenital central hypoventilation syndrome. J Appl Physiol (1985), 116, 439–50.

Corcoran, A. E., Hodges, M. R., Wu, Y., Wang, W., Wylie, C. J., Deneris, E. S. & Richerson, G. B. 2009. Medullary serotonin neurons and central CO2 chemoreception. Respir Physiol Neurobiol, 168, 49–58.

Danneman, P. J. & Mandrell, T. D. 1997. Evaluation of five agents/methods for anesthesia of neonatal rats. Lab Anim Sci, 47, 386–95.

Depuy, S. D., Kanbar, R., Coates, M. B., Stornetta, R. L. & Guyenet, P. G. 2011. Control of breathing by raphe obscurus serotonergic neurons in mice. J Neurosci, 31, 1981–90.

Di Lascio, S., Benfante, R., Cardani, S. & Fornasari, D. 2020. Research Advances on Therapeutic Approaches to Congenital Central Hypoventilation Syndrome (CCHS). Front Neurosci, 14, 615666.

Di Pasquale, E., Monteau, R. & Hilaire, G. 1994. Endogenous serotonin modulates the fetal respiratory rhythm: an in vitro study in the rat. Brain Res Dev Brain Res, 80, 222–32.

Dubreuil, V., Ramanantsoa, N., Trochet, D., Vaubourg, V., Amiel, J., Gallego, J., Brunet, J. F. & Goridis, C. 2008. A human mutation in Phox2b causes lack of CO2 chemosensitivity, fatal central apnea, and specific loss of parafacial neurons. Proc Natl Acad Sci U S A, 105, 1067–1072.

Eddahibi, S., Chaouat, A., Morrell, N., Fadel, E., Fuhrman, C., Bugnet, A. S., Dartevelle, P., Housset, B., Hamon, M., Weitzenblum, E. & Adnot, S. 2003. Polymorphism of the serotonin transporter gene and pulmonary hypertension in chronic obstructive pulmonary disease. Circulation, 108, 1839–44.

Gestreau, C., Heitzmann, D., Thomas, J., Dubreuil, V., Bandulik, S., Reichold, M., Bendahhou, S., Pierson, P., Sterner, C., Peyronnet-Roux, J., Benfriha, C., Tegtmeier, I., Ehnes, H., Georgieff, M., Lesage, F., Brunet, J. F., Goridis, C., Warth, R. & Barhanin, J. 2010. Task2 potassium channels set central respiratory CO2 and O2 sensitivity. Proc Natl Acad Sci U S A, 107, 2325–2330.

Guyenet, P. G. & Bayliss, D. A. 2015. Neural Control of Breathing and CO2 Homeostasis. Neuron, 87, 946–61.

Harper, R. M., Kumar, R., Macey, P. M., Woo, M. A. & Ogren, J. A. 2014. Affective brain areas and sleep-disordered breathing. Prog Brain Res, 209, 275–93.

Hawkins, V. E., Hawryluk, J. M., Takakura, A. C., Tzingounis, A. V., Moreira, T. S. & Mulkey, D. K. 2015. HCN channels contribute to serotonergic modulation of ventral surface chemosensitive neurons and respiratory activity. J Neurophysiol, 113, 1195–205.

Heils, A., Teufel, A., Petri, S., Stober, G., Riederer, P., Bengel, D. & Lesch, K. P. 1996. Allelic variation of human serotonin transporter gene expression. J Neurochem, 66, 2621–4.

Hodges, M. R. & Richerson, G. B. 2008. Contributions of 5-HT neurons to respiratory control: neuromodulatory and trophic effects. Respir Physiol Neurobiol, 164, 222–32.

Holtman, J. R., Jr., Anastasi, N. C., Norman, W. P. & Dretchen, K. L. 1986. Effect of electrical and chemical stimulation of the raphe obscurus on phrenic nerve activity in the cat. Brain Res, 362, 214–20.

Husch, A., Van Patten, G. N., Hong, D. N., Scaperotti, M. M., Cramer, N. & Harris-Warrick, R. M. 2012. Spinal cord injury induces serotonin supersensitivity without increasing intrinsic excitability of mouse V2a interneurons. J Neurosci, 32, 13145–54.

Joubert, F., Loiseau, C., Perrin-Terrin, A. S., Cayetanot, F., Frugiere, A., Voituron, N. & Bodineau, L. 2016a. Key Brainstem Structures Activated during Hypoxic Exposure in One-day-old Mice Highlight Characteristics for Modeling Breathing Network in Premature Infants. Front Physiol, 7, 609.

Joubert, F., Perrin-Terrin, A. S., Verkaeren, E., Cardot, P., Fiamma, M. N., Frugiere, A., Rivals, I., Similowski, T., Straus, C. & Bodineau, L. 2016b. Desogestrel enhances ventilation in ondine patients: Animal data involving serotoninergic systems. Neuropharmacology, 107, 339–350.

Kumar, N. N., Velic, A., Soliz, J., Shi, Y., Li, K., Wang, S., Weaver, J. L., Sen, J., Abbott, S. B., Lazarenko, R. M., Ludwig, M. G., Perez-Reyes, E., Mohebbi, N., Bettoni, C., Gassmann, M., Suply, T., Seuwen, K., Guyenet, P. G., Wagner, C. A. & Bayliss, D. A. 2015. Regulation of breathing by CO(2) requires the proton-activated receptor GPR4 in retrotrapezoid nucleus neurons. Science, 348, 1255–60.

Li, D. C. C.I.H; Kato, R.; Ward, S.L.D; Keens, T.G. Does Desogestrel Improve Ventilatory Control In Congenital Central Hypoventilation Syndrome? ATS 2013, 2013 Philadelphia. ATS society.

Loiseau, C., Casciato, A., Barka, B., Cayetanot, F. & Bodineau, L. 2019. Orexin Neurons Contribute to Central Modulation of Respiratory Drive by Progestins on ex vivo Newborn Rodent Preparations. Front Physiol, 10, 1200.

Loiseau, C., Cayetanot, F., Joubert, F., Perrin-Terrin, A. S., Cardot, P., Fiamma, M. N., Frugiere, A., Straus, C. & Bodineau, L. 2018. Current Perspectives for the use of Gonane Progesteronergic Drugs in the Treatment of Central Hypoventilation Syndromes. Curr Neuropharmacol, 16, 1433–1454.

Loiseau, C., Osinski, D., Joubert, F., Straus, C., Similowski, T. & Bodineau, L. 2014. The progestin etonogestrel enhances the respiratory response to metabolic acidosis in newborn rats. Evidence for a mechanism involving supramedullary structures. Neurosci Lett, 567, 63–7.

Menuet, C., Kourdougli, N., Hilaire, G. & Voituron, N. 2011. Differences in serotoninergic metabolism possibly contribute to differences in breathing phenotype of FVB/N and C57BL/6J mice. J Appl Physiol (1985), 110, 1572–81.

Morin, D., Monteau, R. & Hilaire, G. 1991. Serotonin and cervical respiratory motoneurones: intracellular study in the newborn rat brainstem-spinal cord preparation. Exp Brain Res, 84, 229–32.

Morin, L. P., Michels, K. M., Smale, L. & Moore, R. Y. 1990. Serotonin regulation of circadian rhythmicity. Ann N Y Acad Sci, 600, 418–26.

Murakoshi, T., Suzue, T. & Tamai, S. 1985. A pharmacological study on respiratory rhythm in the isolated brainstem-spinal cord preparation of the newborn rat. Br J Pharmacol, 86, 95–104.

Niblock, M. M., Lohr, K. M., Nixon, M., Barnes, C., Schaudies, M. & Murphy, M. 2012. Cells in the female retrotrapezoid region upregulate c-fos in response to 10%, but not 5%, carbon dioxide. Brain Res, 1433, 62–8.

Onimaru, H. & Homma, I. 2003. A novel functional neuron group for respiratory rhythm generation in the ventral medulla. J Neurosci, 23, 1478–1486.

Onimaru, H., Ikeda, K. & Kawakami, K. 2008. CO2-sensitive preinspiratory neurons of the parafacial respiratory group express Phox2b in the neonatal rat. J Neurosci, 28, 12845–50.

Onimaru, H., Ikeda, K., Mariho, T. & Kawakami, K. 2014. Cytoarchitecture and CO(2) sensitivity of Phox2b-positive Parafacial neurons in the newborn rat medulla. Prog Brain Res, 209, 57–71.

Paxinos, G. & Franklin, K. B. 2001. The mouse brain in stereotaxic coordinates, 2nd Edition, San Diego, CA: Academic Press.

Paxinos, G., Halliday, G., Watson, C., Koutcherov, Y. & Wang, H. 2007. Atlas of the developping mouse brain, Amsterdam, Academic Press.

Peever, J. H., Necakov, A. & Duffin, J. 2001. Nucleus raphe obscurus modulates hypoglossal output of neonatal rat in vitro transverse brain stem slices. J Appl Physiol (1985), 90, 269–79.

Ramanantsoa, N., Hirsch, M. R., Thoby-Brisson, M., Dubreuil, V., Bouvier, J., Ruffault, P. L., Matrot, B., Fortin, G., Brunet, J. F., Gallego, J. & Goridis, C. 2011. Breathing without CO(2) chemosensitivity in conditional Phox2b mutants. J Neurosci, 31, 12880–8.

Ramirez, J. M. & Viemari, J. C. 2005. Determinants of inspiratory activity. Respir Physiol Neurobiol, 147, 145–57.

Ren, J. & Greer, J. J. 2006. Neurosteroid modulation of respiratory rhythm in rats during the perinatal period. J.Physiol, 574, 535–546.

Richerson, G. B. 2004. Serotonergic neurons as carbon dioxide sensors that maintain pH homeostasis. Nat Rev Neurosci, 5, 449–61.

Song, N., Zhang, G., Geng, W., Liu, Z., Jin, W., Li, L., Cao, Y., Zhu, D., Yu, J. & Shen, L. 2012. Acid sensing ion channel 1 in lateral hypothalamus contributes to breathing control. PLoS One., 7, e39982.

Sritippayawan, S., Hamutcu, R., Kun, S. S., Ner, Z., Ponce, M. & Keens, T. G. 2002. Mother-daughter transmission of congenital central hypoventilation syndrome. Am J Respir Crit Care Med, 166, 367–9.

Straus, C., Trang, H., Becquemin, M. H., Touraine, P. & Similowski, T. 2010. Chemosensitivity recovery in Ondine’s curse syndrome under treatment with desogestrel. Respir Physiol Neurobiol, 171, 171–4.

Suzue, T. 1984. Respiratory rhythm generation in the in vitro brain stem-spinal cord preparation of the neonatal rat. J Physiol, 354, 173–83.

Trang, H., Samuels, M., Ceccherini, I., Frerick, M., Garcia-Teresa, M. A., Peters, J., Schoeber, J., Migdal, M., Markstrom, A., Ottonello, G., Piumelli, R., Estevao, M. H., Senecic-Cala, I., Gnidovec-Strazisar, B., Pfleger, A., Porto-Abal, R. & Katz-Salamon, M. 2020. Guidelines for diagnosis and management of congenital central hypoventilation syndrome. Orphanet J Rare Dis, 15, 252.

Veasey, S. C., Fornal, C. A., Metzler, C. W. & Jacobs, B. L. 1995. Response of serotonergic caudal raphe neurons in relation to specific motor activities in freely moving cats. J Neurosci, 15, 5346–59.

Voiculescu, O., Charnay, P. & Schneider-Maunoury, S. 2000. Expression pattern of a Krox-20/Cre knock-in allele in the developing hindbrain, bones, and peripheral nervous system. Genesis, 26, 123–6.

Voituron, N., Frugiere, A., Champagnat, J. & Bodineau, L. 2006. Hypoxia-sensing properties of the newborn rat ventral medullary surface in vitro. J Physiol, 577, 55–68.

Voituron, N., Frugiere, A., Mc Kay, L. C., Romero-Granados, R., Dominguez-Del-Toro, E., Saadani-Makki, F., Champagnat, J. & Bodineau, L. 2011. The kreisler mutation leads to the loss of intrinsically hypoxia-activated spots in the region of the retrotrapezoid nucleus/parafacial respiratory group. Neuroscience, 194, 95–111.

Voituron, N., Shvarev, Y., Menuet, C., Bevengut, M., Fasano, C., Vigneault, E., El Mestikawy, S. & Hilaire, G. 2010. Fluoxetine treatment abolishes the in vitro respiratory response to acidosis in neonatal mice. PLoS One, 5, e13644.

Weese-Mayer, D. E., Berry-Kravis, E. M., Ceccherini, I., Keens, T. G., Loghmanee, D. A., Trang, H. & Subcommittee, A. T. S. C. C. H. S. 2010. An official ATS clinical policy statement: Congenital central hypoventilation syndrome: genetic basis, diagnosis, and management. Am J Respir Crit Care Med, 181, 626–44.

Weese-Mayer, D. E., Rand, C. M., Zhou, A., Carroll, M. S. & Hunt, C. E. 2017. Congenital central hypoventilation syndrome: a bedside-to-bench success story for advancing early diagnosis and treatment and improved survival and quality of life. Pediatr Res, 81, 192–201.

Weese-Mayer, D. E., Silvestri, J. M., Menzies, L. J., Morrow-Kenny, A. S., Hunt, C. E. & Hauptman, S. A. 1992. Congenital central hypoventilation syndrome: diagnosis, management, and long-term outcome in thirty-two children. J Pediatr, 120, 381–7.

Wei, A. D. & Ramirez, J. M. 2019. Presynaptic Mechanisms and KCNQ Potassium Channels Modulate Opioid Depression of Respiratory Drive. Front Physiol, 10, 1407.

